# The Quagmire of Immunogenicity-Guided Antibody Design: Labels Limit Progress

**DOI:** 10.64898/2026.10.27.762175

**Authors:** Amelia Villegas-Morcillo, Chirag Raman, Jana M. Weber, Marcel J.T. Reinders

## Abstract

Generative sequence-based antibody design is increasingly steered by learned property predictors, so a design objective is only as well specified as the labels behind its oracle. Immunogenicity is among the most clinically consequential properties to condition on, and among the least well specified: the supervisory signal is a single anti-drug-antibody percentage (ADA%) per molecule, aggregated from published trials. To analyze that signal, we first extended and corrected the widely used 217-entry dataset to 231 therapeutics with paired sequences, spanning formats from mouse to fully human. Then, we manually reconstructed the clinical evidence for 110 of them from primary sources. We find that an antibody format lookup outperforms every sequence-based learned predictor on the 231-antibody dataset. Apparent accuracy tracks the format composition of the evaluation split rather than sequence signal. Furthermore, the labels understate their own evidence: molecules labeled at *≤* 6 ADA% span up to 86% across trials. We recomputed labels from the trial-level measurements and found that the advantage of the format-based approach disappears, while the sequence-based predictors degrade less under the cohort-weighted mean. We conclude that progress is limited by the label itself, and recommend reporting ADA% at the trial level, treating the target as a distribution rather than a point value, and evaluating against a format-based baseline.

## 1 Introduction

Generative models can now propose antibody sequences and structures that plausibly bind a specified target [1]. Binding, however, is a necessary but not a sufficient condition for a therapeutic: before it can reach the clinic, a candidate must also be expressible, thermostable, and safe in patients, who can develop an antibody response against the therapeutic itself [2, 3]. These developability and safety constraints enter a generative pipeline through learned property predictors: as conditioning variables, rewards, acquisition targets in multi-objective optimization [4], or tools called by an LLM-driven design agent [5]. A generative model is therefore as well specified as the oracle it optimizes against, and that oracle is only as well specified as its labels.

This paper is a study of reported labels in accessible databases. Our original aim was a conditional generative model for immunogenicity-risk reduction, for which the prerequisite was an immunogenicity oracle. Building that oracle taught us what makes the task difficult, and we show that this is the reported data rather than the model. The experiments here diagnose sequence-based immunogenicity prediction, not a design loop. Our conclusions nonetheless apply to any broader pipeline that uses immunogenicity prediction as guiding signal.

### Immunogenicity is a hard supervision problem

Immunogenicity is measured clinically as the percentage of treated patients that develop anti-drug antibodies (ADA%), which can accelerate clearance of the therapeutic, reduce efficacy, and cause hypersensitivity and infusion reactions [6, 7]. Two features make this a difficult regression target for machine learning. It is *scarce*: of roughly a thousand therapeutic antibodies in public databases [8, 9], only a few hundred have accessible ADA% information, one value per molecule. And it is *extrinsic*: ADA% is not a measurement of the molecule alone, but the outcome across a cohort of patients. It therefore carries the dose, the route of administration, the indication, the comedication, the assay, and the duration of follow-up along with it; yet none of this is reported.

### Humanness as a proxy

Because ADA% decreases as therapeutic antibody engineering moves from mouse to chimeric, humanized and fully human formats, *humanness* became the dominant computational proxy for immunogenicity risk [10]. Sequence-based scores quantify resemblance to natural human repertoires [11–14], and are in turn used as optimization objectives for humanization and design [15–18]. This is a desirable proxy because humanness is a well-defined function of the protein sequence. However, even fully human antibodies can still elicit a wide range of ADA%, and humanization alone does not determine which molecules will be immunogenic [19]. Parkinson and Wang [20] make this concrete: humanness scores track ADA% across antibody formats but not within them. Mechanism-based alternatives predict MHC class II peptide presentation [21], since ADA formation requires peptides from the therapeutic to be presented to CD4^+^ T cells. A further line of research trains directly on curated ADA% [22–25]. All of these rely on the same supervision signal: one ADA% value per antibody as reported.

### Contributions

Recent large-scale studies have established that humanness metrics and embedding-based models correlate weakly with clinical ADA% [26, 27], a weakness attributed mainly to the predictors, though Dudzic and Krawczyk [26] also point to clinical metadata and data quality. We pursue that second explanation, analyzing the supervision signal itself with a dataset whose clinical origin we curate ourselves.

- We correct and extend the 217-entry Hu-mAb [11] dataset to **231** therapeutics with paired heavy and light chains, and manually curate the underlying clinical evidence for **110** of them (426 records) from regulatory documents, trial registries, and primary literature (Section 2).
- We show that a lookup table returning the average ADA% per antibody format in each training set outperforms every learned predictor on this dataset, indicating that per-fold accuracy tracks the format composition of the evaluation split rather than sequence signal (Section 3.1).
- We show that the reported ADA% labels do not reflect the evidence behind them: we found therapeutics labeled at *≤*6% to have individual trials reporting as high as 86% (Section 3.2).
- We show that recomputing curated labels as a cohort-weighted mean reduces the performance of the format-based approach more than that of the sequence-based predictors. This indicates that accuracy is partly an artifact of how ADA% is summarized across trials (Section 3.3).
- We set out guidelines for the community to make a step forward in building immunogenicity-guided generative models (Section 4).

## 2 Data and models

### From 217 to 231 therapeutics

Most published immunogenicity predictors train or validate against the 217 entries from Hu-mAb [11]. Analyzing that dataset, we found it covers 214 distinct molecules: three entries are the same molecule listed twice under different names, each carrying a different ADA% value from a different reference (Afutuzumab/Obinutuzumab 6.6% vs. 6.0%; PankoMab/Gatipotuzumab 25.0% vs. 9.5%; Ticilimumab/Tremelimumab 2.0% vs. 0.0%). These duplications are an early reflection of this paper’s argument: one molecule, but two names with two ground truths inside the original dataset. We retain one entry per molecule, add 17 antibodies approved between 2019 and 2023 from [28], and cross-check every sequence against Thera-SAbDab [8]. This results in **231** therapeutics with paired chains, format annotation, and an ADA%; 97 additionally have a determined structure. The format annotation records how much of each antibody sequence is derived from human versus non-human (typically mouse) origin: mouse, chimeric, humanized, and fully human. The dataset is dominated by human and humanized molecules, but mouse and chimeric carry most of the label variance (Figure 1a; format composition in Appendix A).

**Figure 1:**
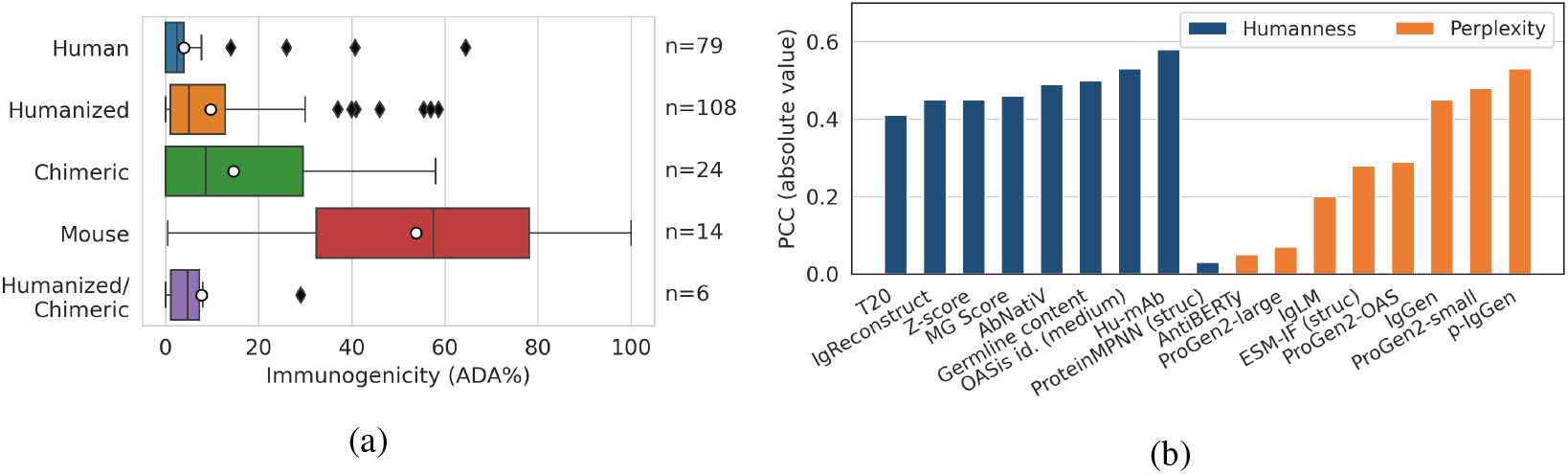
Antibody format distinguishes immunogenicity better than humanness tracks it. Distribution of ADA% by antibody format across the 231 therapeutics, with format counts. Pearson correlation (PCC) between humanness scores (blue bars) and protein language-model perplexity (orange bars) against ADA% for the original 217-entry dataset. No score exceeds 0.6.

### Reconstructing the clinical evidence for 110 therapeutics

The reported ADA% in the original dataset is a single number whose full derivation is not recorded. To recover it, we searched each antibody by name across FDA and EMA regulatory documents, ClinicalTrials.gov, and primary literature, and transcribed every reported ADA% measurement together with its clinical context, including cohort size, route, and indication (protocol and recorded fields in Appendix B). This produced **426** trial-level records for **110** therapeutics, a median of 4 records each and up to 16.

### Models

Heavy and light chains are embedded with AntiBERTy [29], mean-pooled per chain and averaged (512-dim). We use amino-acid composition (20-dim) as a sequence-statistics control. Predictors are a 2-layer MLP regressor (100–50, ReLU) trained with MSE, and Rank-N-Contrast (RNC) [30], a supervised contrastive objective for continuous targets that orders representations by label distance. Evaluation is 5-fold cross-validation, with folds balanced on four ADA% ranges (Appendix C). Using the same CV splits, we repeat this 10 times to observe variance in the runs. We pool predictions from all test folds and compute metrics against the full set of 231 labels. We report mean absolute error (MAE), Pearson correlation (PCC), and *R*^2^. As baselines, we also compare to zero-shot antibody language-model perplexity (AntiBERTy and p-IgGen [31]), a label-free measure of how likely the model finds each sequence; and the average ADA% per antibody format (mouse, chimeric, humanized/chimeric, humanized, and human; 5 values in each CV split), which we refer to as the format-based lookup approach (details in Appendix D).

## 3 Results

### 3.1 Benchmark accuracy tracks format composition, not sequence

Antibody format and humanness are properties of the molecule alone, but ADA% is a property of the molecule in a specific treated cohort. Figure 1a shows that, across the 231 therapeutics, format separates the label strongly. This relationship is well known; here we quantify the predictive power of format alone. Mouse therapeutics have an average ADA% of 53.78 against 3.96 for fully human ones, while no humanness or perplexity score correlates above 0.6 with ADA% within the 217-entry dataset (Figure 1b; scores are taken from [12, 31]).

When evaluating in a cross-validation setting, we pooled the predictions of the 5 test folds and compared them to the 231 original ADA% values. Figure 2a shows that the format-based approach reaches MAE=9.4, PCC=0.58, *R*^2^=0.33, outperforming every learned predictor. AntiBERTy embeddings with an MLP reach PCC=0.47, *R*^2^=0.20 and zero-shot p-IgGen perplexity PCC=0.49 without any training at all. The amino-acid composition control reaches only PCC=0.29, so neither sequence statistics nor learned embeddings match the format-based lookup. Note that the p-IgGen perplexity correlation reported here is slightly lower than in Figure 1b, since it is recomputed over the extended 231-antibody dataset rather than the original 217. Inspecting the learned representations confirms the impact of the antibody format. The original AntiBERTy embedding space (Figure 2b) is already organized by format before any supervision. RNC reshapes this space into a smooth one-dimensional ordering by label but performs worse than the MLP, even after adding an MSE term (PCC=0.32, *R*^2^=0.08).

**Figure 2:**
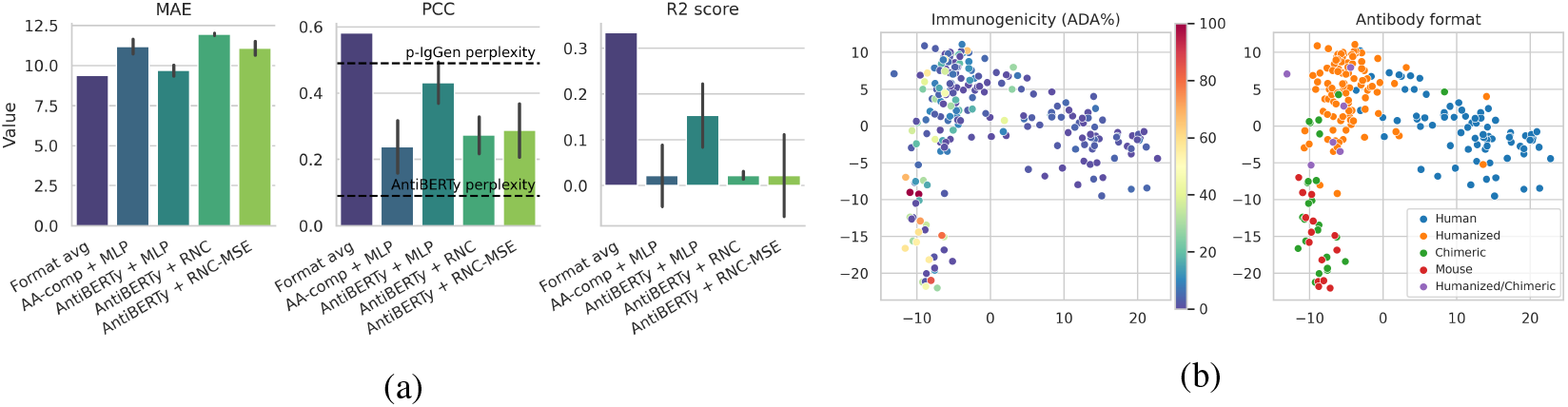
Apparent accuracy is a property of the benchmark, not of the model. (a) Held-out MAE, PCC and *R*^2^ on the 231-therapeutic dataset (5-folds pooled, mean *±* SD over 10 runs, dashed lines are zero-shot language-model perplexity baselines); the format-based lookup outperforms every learned predictor. (b) PCA of AntiBERTy embeddings colored by ADA% (left) and by antibody format (right): the space is mostly organized by format before any supervision.

Performance also varies sharply across CV folds (Figure 8, Appendix E). Because folds are balanced on ADA% alone, format composition can differ vastly between them. For example, test folds 4 and 5 contain no mouse antibodies, so they evaluate humanized and chimeric molecules against each other. For these folds, *R*^2^ falls to or below zero for every model, including the format-based approach.

### 3.2 The label is a distribution reported as a number

The 110 curated therapeutics let us compare each original ADA% value against the trial-level measurements it summarizes. Figure 3 shows 67 molecules labeled at *≤* 6% in the original dataset, which are those a design pipeline would treat as safe. For each, we plot every trial-level measurement we recovered, together with the number of records we found.

**Figure 3:**
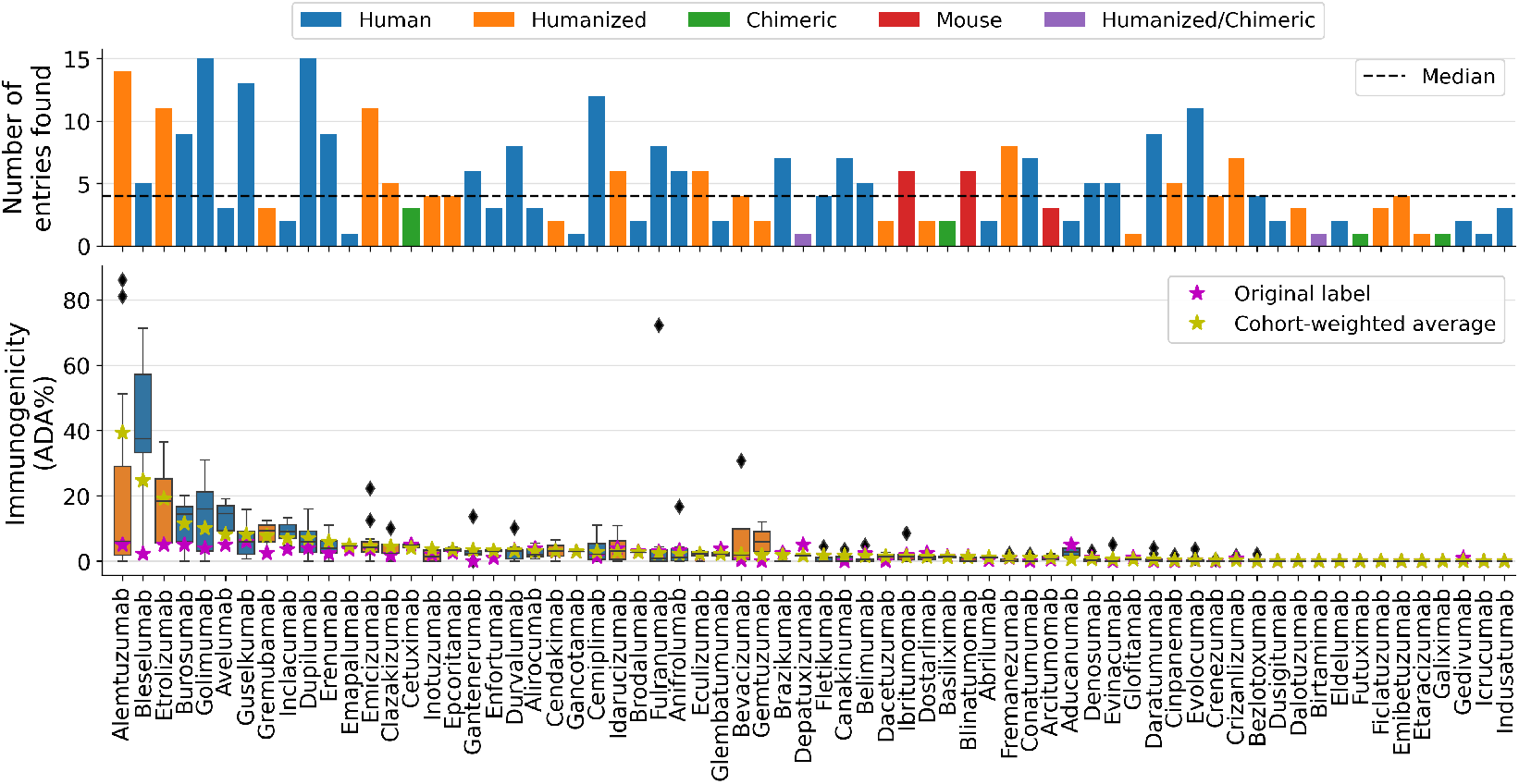
Reported ADA% measurements for therapeutics in the original dataset vary across trials. Shown for the 67 of the 110 curated therapeutics whose original ADA% value is *≤*6%, and colored by antibody format. Top: number of trial-level records found per therapeutic. Bottom: each box is one antibody’s ADA% across the trials we curated; the magenta star is the original dataset value and the yellow star the cohort-weighted average recomputed from the same evidence.

Figure 3 shows that the original reported ADA% is a poor summary of the measurements behind it. Alemtuzumab is reported at ~ 5% but spans a range of 0–86% across 14 trials, with a cohort-weighted mean near 39%. Bevacizumab spans 0–30.7%, burosumab 0–20%, cemiplimab 0–11.1%. For these 67 therapeutics, the median within-antibody spread is 4.8 percentage points (IQR 2.0–14.0) and the mean is 12.3, so a minority of molecules carry very wide spreads. Part of this spread is expected: cohorts can be small, and a proportion measured in a few dozen patients is uncertain. But the ranges are far wider than their cohort sizes account for, and a model trained on a single ADA% must absorb that variation as noise.

### 3.3 Recomputed labels from trials influence prediction performance

We recomputed labels for the 110 curated therapeutics using two alternatives: the cohort-weighted mean, which estimates the expected ADA% across different-sized populations, and the maximum observed value, which is the quantity relevant for a safety-screening pipeline.

Figures 4 and 9 (per-fold in Figure 10, Appendix E) show that performance drops for both re-labelings using this smaller dataset of 110 therapeutics. Using the original reported labels, the format-based approach is still the strongest predictor (PCC=0.52, *R*^2^=0.26), ahead of AntiBERTy with MLP or RNC-MSE. When applying the cohort-weighted mean, the ordering inverts: the format-based lookup collapses to PCC=0.34, *R*^2^=0.03, while the MLP model retains PCC=0.43, *R*^2^=0.17. Zero-shot p-IgGen perplexity follows the format-based approach down, from PCC=0.53 to 0.45 and 0.34. Under the maximum label, the sequence models fall further than the format-based lookup, with AntiBERTy + RNC-MSE dropping from PCC=0.38 to 0.10.

**Figure 4:**
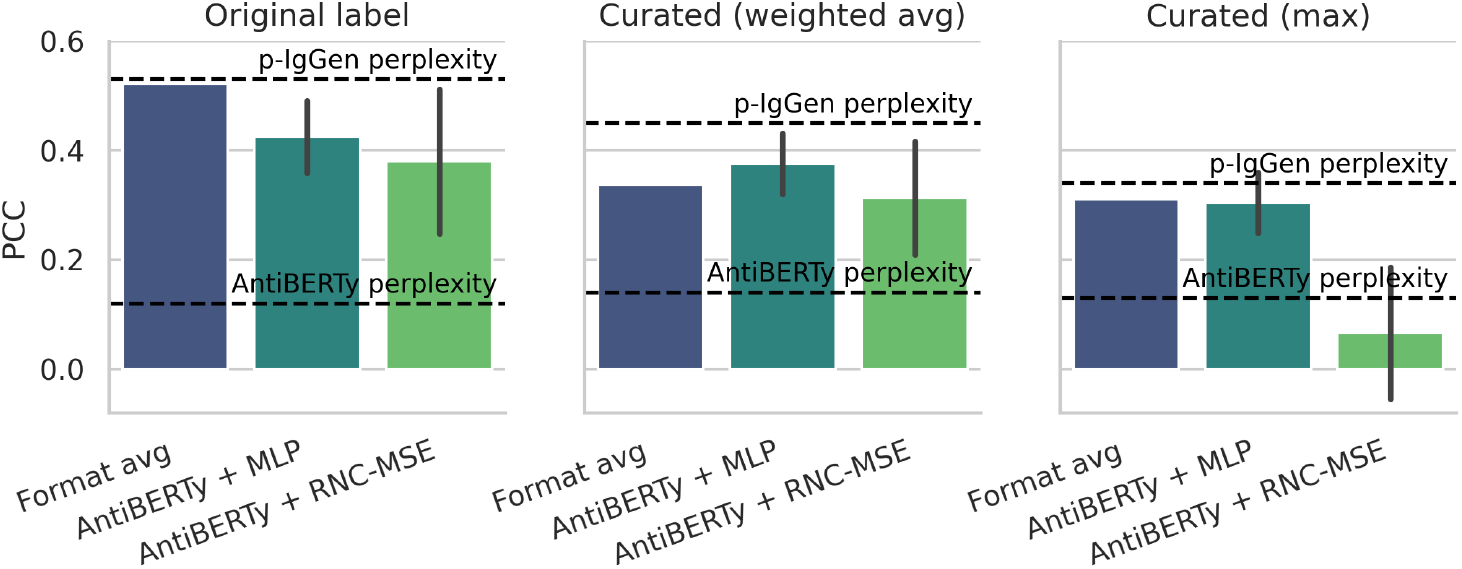
Recomputed ADA% labels influence prediction performance. Held-out PCC on 110 curated therapeutics under the original label (left), the cohort-weighted mean (middle), and the maximum observed value (right). Both curated labels reduce the format-based lookup performance, while AntiBERTy + RNC-MSE drops mainly under the maximum. 5-folds pooled, mean *±* SD over 10 runs, dashed lines are zero-shot language-model perplexity. MAE and *R*^2^ in Figure 9.

Under the cohort-weighted mean, the format-based approach degrades far more than the sequence models do. The original label is a single reported value with unclear derivation, often from one source, while the curated ones are computed from every trial we found. The curated labels shift unevenly across formats (Table 2, Appendix A): the human and humanized means increase while the mouse and chimeric means barely change. This brings the format means closer together, most strongly under the maximum label, reducing the separation the format-based approach depends on. Moreover, the two curated labels behave differently: the cohort-weighted mean stabilizes while the maximum can only rise with more records. Under the maximum, the within-format variation partly reflects how much evidence we found rather than the molecule.

**Table 1:** Format composition of the extended dataset with 231 therapeutic antibodies (SD: standard deviation, MAD: median absolute deviation). For most formats (except mouse), MAD is comparable to the median, reflecting a zero-inflated, right-skewed distribution (as seen in Figure 1a).

| Antibody format | Count | Mean ± SD ADA% | Median (MAD) ADA% |
|---|---|---|---|
| Human | 79 | 3.96 ± 8.87 | 2.4 (2.0) |
| Humanized | 108 | 9.70 ± 12.87 | 5.0 (4.5) |
| Chimeric | 24 | 14.55 ± 16.35 | 8.6 (8.6) |
| Mouse | 14 | 53.78 ± 34.56 | 57.5 (25.0) |
| Humanized/chimeric | 6 | 7.75 ± 10.86 | 4.75 (4.0) |
| Total | 231 | 10.86 ± 18.20 | 3.5 (3.5) |

**Table 2:** Mean ±SD ADA% per antibody format for the 110 curated therapeutics, using the original label and the two recomputed curated labels.

| Antibody format | Count | Original label | Curated |  |
|---|---|---|---|---|
|  |  |  | Cohort-weighted mean | Maximum |
| Human | 40 | 4.2 ± 10.2 | 5.3 ± 10.6 | 12.3 ± 18.9 |
| Humanized | 46 | 8.8 ± 11.2 | 11.1 ± 13.3 | 15.9 ± 18.4 |
| Chimeric | 14 | 21.8 ± 17.5 | 19.0 ± 16.6 | 22.1 ± 18.3 |
| Mouse | 8 | 47.5 ± 44.4 | 46.9 ± 44.1 | 48.8 ± 43.1 |
| Humanized/chimeric | 2 | 2.5 ± 3.5 | 0.8 ± 1.2 | 0.8 ± 1.2 |
| Total | 110 | 11.5 ± 19.6 | 12.4 ± 19.6 | 17.5 ± 22.8 |

The smaller drop for the sequence models may indicate format-independent sequence signal. However, the AntiBERTy space is already organized by format before any supervision (Figure 2b), so a sequence model can track the differences between formats without resolving variation inside them. After removing format, the format-based approach carries no within-format information by construction, so its correlation falls from PCC = 0.52 to zero. In contrast, AntiBERTy + MLP retains PCC = 0.13 [ *−*0.07, 0.32] under the original labels and 0.13 [ *−*0.06, 0.31] under the cohort-weighted mean (Table 6, Appendix E). The sequence models therefore keep some information once format is removed, but it is small and the confidence intervals include zero.

## 4 Discussion

### What is needed

Three changes would make immunogenicity a better-posed target for generative models. The first is to publish the trial-level information together with any curated ADA%. These are reported in regulatory documents and usually discarded during curation. The second is to treat the target as a distribution rather than a point. The output of a sequence model should then be a wide, calibrated interval, and a design loop could act on this. The third is to report the format-based baseline. Because antibody format explains much of the label variance in legacy datasets, reported accuracy should be compared against what this baseline can already achieve.

### Convergent evidence

Two concurrent studies reach compatible conclusions from different directions: Dudzic and Krawczyk [26] assemble a therapeutic antibody database from trial registries and find that humanness metrics do not predict immunogenicity, and Chungyoun and Gray [27] report weak correlations for embedding-based predictors on a larger developability benchmark. Agnihotri et al. [32] independently argue for shared immunogenicity data infrastructure. This convergence suggests that the limitation is not the modeling choice. Our contribution is orthogonal: by reconstructing the clinical evidence beneath the labels, we show that format composition inflates apparent performance, and that recomputing labels removes this shortcut while affecting the sequence-based predictors less.

### Limitations

Our trial-level curation covers 110 therapeutics and is not exhaustive. For the antibodies it covers, we cannot ensure every trial was found, so the observed spread is a lower bound on the true heterogeneity. We also fix a single embedding model, since different options have been previously analyzed [27], and our conclusions concern the label’s reliability, not the choice of representation.

## A Extended dataset details

## B Curation protocol

We searched each antibody name in: (i) Food and Drug Administration (FDA) approved drugs (https://www.accessdata.fda.gov/scripts/cder/daf/index.cfm); (ii) European Medicines Agency (EMA) medicines (https://www.ema.europa.eu/en/medicines); (iii) clinicaltrials.gov results; and (iv) primary literature.

For each found trial we recorded: trial identifier or study name, indication, route of administration, dose, number of exposures, follow-up window at which ADA was assessed, cohort size, percentage of patients with ADA (ADA%), assay type (ELISA, ECL, EIA, drug-tolerant EIA), antibody type (HAMA, HACA, HAHA, AIA), and source URL. Where a source reported an ADA% range rather than a point value, we recorded the range and used its midpoint. We retained the original label for those antibodies for which we could not find any primary source reporting ADA% measurements.

## C Cross-validation splits and fold composition

Folds are balanced on the label using four ADA% ranges, whose boundaries are recomputed per label definition so that the range groups remain equally populated (Table 3). Since balancing is applied to the label alone, format composition varies between folds (Figure 5). Test folds 1–3 retain 6–11% mouse antibodies, while test folds 4 and 5 contain no mouse antibodies at all. Test fold 3 most directly contrasts human against mouse molecules. The differences in performance across folds are therefore a property of the evaluation split. To control for variance introduced by the evaluation split itself, we recommend balancing splits jointly on ADA% label and format. This complements the format-based baseline, which tests whether accuracy reflects format alone or genuine sequence signal, making it the more direct diagnostic and the one we recommend first.

**Table 3:** Range boundaries (ADA%) used to balance the cross-validation splits.

| Dataset | Label definition | Range boundaries |  |  |  |
|---|---|---|---|---|---|
| Extended 231 antibodies | Original label | [0, 0.6] | (0.6, 3.6] | (3.6, 11.9] | (11.9, 100] |
| Curated 110 antibodies | Original label | [0, 1.0] | (1.0, 3.6] | (3.6, 9.5] | (9.5, 100] |
|  | Curated (cohort-weighted average) | [0, 1.3] | (1.3, 4.6] | (4.6, 15.1] | (15.1, 100] |
|  | Curated (maximum) | [0, 2.8] | (2.8, 9.0] | (9.0, 25.1] | (25.1, 100] |

**Figure 5:**
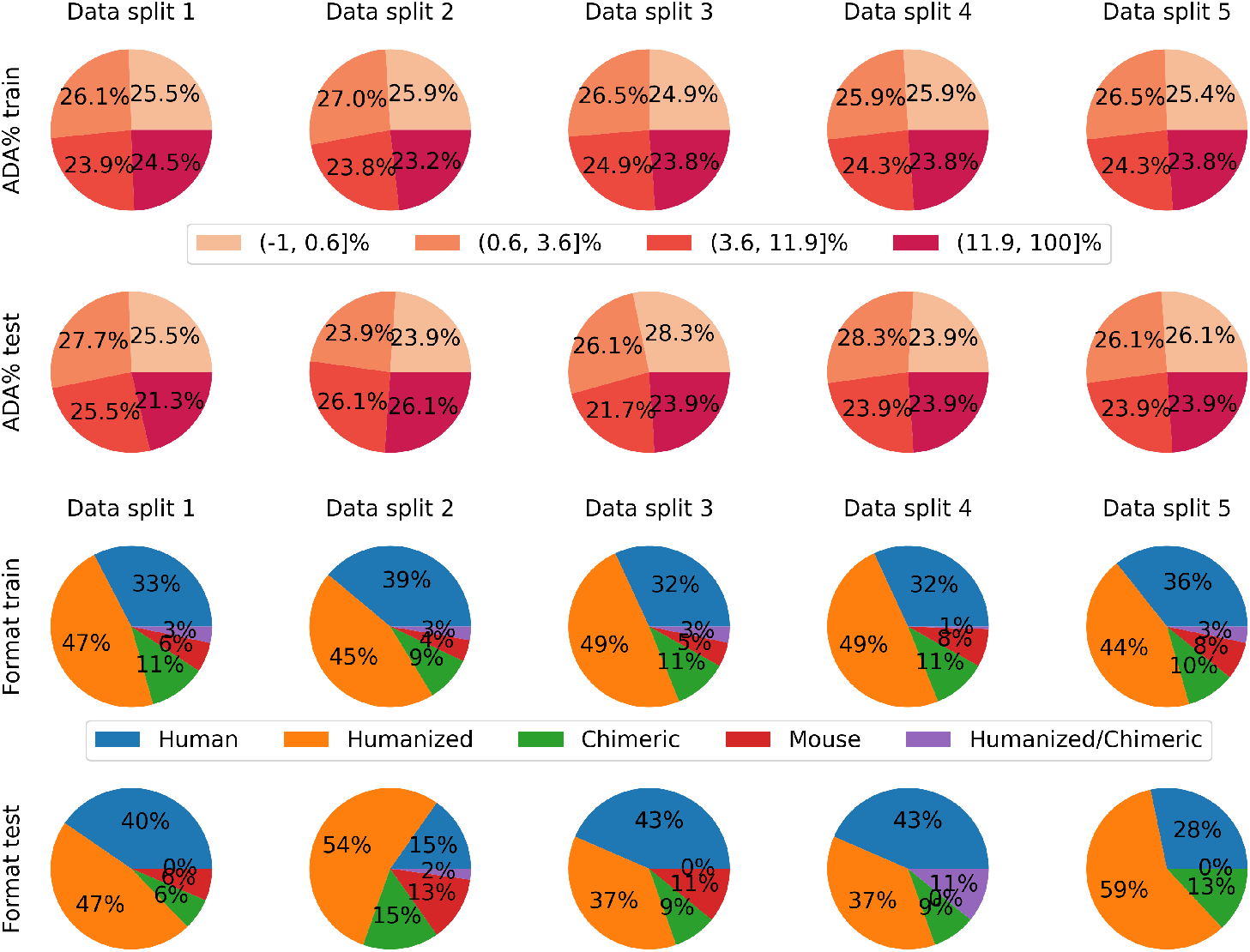
Label composition (top two rows) and antibody format composition (bottom two rows) of the training and test splits of each cross-validation fold. Labels are balanced by ADA% but not format composition, so mouse antibodies are absent from the test folds 4 and 5.

## D Format-based approach

For each of the 5 cross-validation folds, we compute the average ADA% per antibody format on only that fold’s training set. Then, we use these averages to predict the held-out test set: an antibody is assigned the mean ADA% of its own format among the training antibodies. This is repeated independently for every fold, so each fold’s format averages are computed without access to that fold’s test labels. Predictions from all 5 folds are then pooled before computing MAE, PCC, and *R*^2^, consistent with how the learned predictors are evaluated.

## E Extended results

Tables 4 and 5 report 5-fold cross-validation results, with predictions pooled across the 5 test folds and averaged over 10 runs using the same CV splits. Table 6 reports PCC within each antibody format, and after removing format, for the 110 therapeutics with curated labels, using the same pooling and averaging. Figures 6 and 7 show the predicted-versus-observed ADA% values from which we compute the MAE, PCC, and *R*^2^ metrics in the tables.

**Table 4:** Performance on the 231-antibody dataset.

| Model | MAE (↓) | PCC (↑) | *R*<sup>2</sup> (↑) |
|---|---|---|---|
| Format-based lookup | **9.4** | **0.58** | **0.33** |
| AA-composition + MLP | 10.8 | 0.29 | 0.08 |
| AntiBERTy + MLP | **9.4** | 0.47 | 0.20 |
| AntiBERTy + RNC | 12.0 | 0.30 | 0.02 |
| AntiBERTy + RNC-MSE | 10.8 | 0.32 | 0.08 |
| AntiBERTy perplexity (zero-shot) | — | 0.09 | — |
| p-IgGen perplexity (zero-shot) | — | 0.49 | — |

**Table 5:**
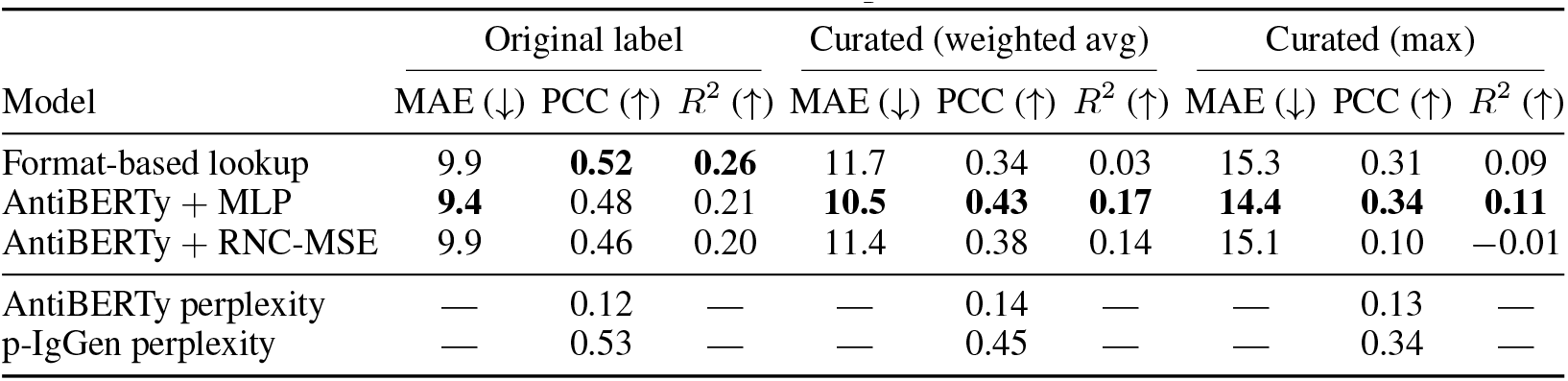
Performance on the 110 curated therapeutics under three label definitions.

|  | Original label |  |  | Curated (weighted avg) |  |  | Curated (max) |  |  |
|---|---|---|---|---|---|---|---|---|---|
| Model | MAE (↓) | PCC (↑) | *R*<sup>2</sup> (↑) | MAE (↓) | PCC (↑) | *R*<sup>2</sup> (↑) | MAE (↓) | PCC (↑) | *R*<sup>2</sup> (↑) |
| Format-based lookup | 9.9 | **0.52** | **0.26** | 11.7 | 0.34 | 0.03 | 15.3 | 0.31 | 0.09 |
| AntiBERTy + MLP | **9.4** | 0.48 | 0.21 | **10.5** | **0.43** | **0.17** | **14.4** | **0.34** | **0.11** |
| AntiBERTy + RNC-MSE | 9.9 | 0.46 | 0.20 | 11.4 | 0.38 | 0.14 | 15.1 | 0.10 | −0.01 |
| AntiBERTy perplexity | — | 0.12 | — | — | 0.14 | — | — | 0.13 | — |
| p-IgGen perplexity | — | 0.53 | — | — | 0.45 | — | — | 0.34 | — |

**Table 6:** Within-format PCC on the 110 curated therapeutics. Partial PCC controls for format and is computed by subtracting each format’s mean from both the label and the prediction before correlating; the 95% CI is a stratified bootstrap (2000 resamples). For humanized/chimeric (count = 2), the correlation is ±1 by construction. We exclude the format-based approach because its prediction is constant within a format.

| Label & Model | PCC<br>(110) | Within-format PCC |  |  |  | Partial PCC [95% CI]<br>(110) |
|---|---|---|---|---|---|---|
|  |  | Human<br>(40) | Humanized<br>(46) | Chimeric<br>(14) | Mouse<br>(8) |  |
| *Original label* |  |  |  |  |  |  |
| AntiBERTy + MLP | 0.48 | 0.32 | 0.13 | 0.31 | −0.07 | +0.13 [−0.07, +0.32] |
| AntiBERTy + RNC-MSE | 0.46 | 0.45 | 0.08 | 0.08 | 0.20 | +0.11 [−0.11, +0.31] |
| *Curated (weighted avg)* |  |  |  |  |  |  |
| AntiBERTy + MLP | 0.43 | 0.39 | 0.13 | 0.30 | −0.14 | +0.13 [−0.06, +0.31] |
| AntiBERTy + RNC-MSE | 0.38 | 0.16 | 0.01 | 0.24 | 0.01 | +0.07 [−0.10, +0.24] |
| *Curated (max)* |  |  |  |  |  |  |
| AntiBERTy + MLP | 0.34 | 0.25 | 0.22 | 0.06 | 0.58 | +0.22 [+0.06, +0.35] |
| AntiBERTy + RNC-MSE | 0.10 | 0.12 | −0.05 | −0.20 | 0.44 | −0.01 [−0.17, +0.15] |

**Figure 6:**
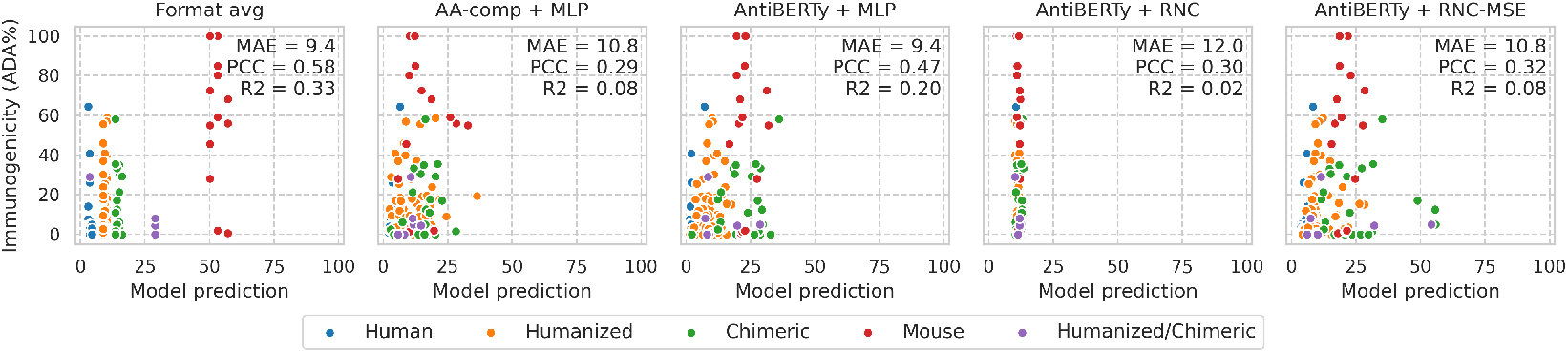
Predicted against observed ADA% on the 231-antibody dataset (5-folds pooled, mean prediction over 10 runs), colored by antibody format. The format-based approach produces one prediction per format category (slightly different in each of the 5 CV splits) and still yields the highest correlation; the learned models spread predictions continuously without separating molecules within a format. Additionally, AntiBERTy + RNC predictions cluster near each training-fold mean, since the contrastive objective accounts only for relative label ordering, not absolute scale; RNC-MSE recovers this by adding a direct regression term.

**Figure 7:**
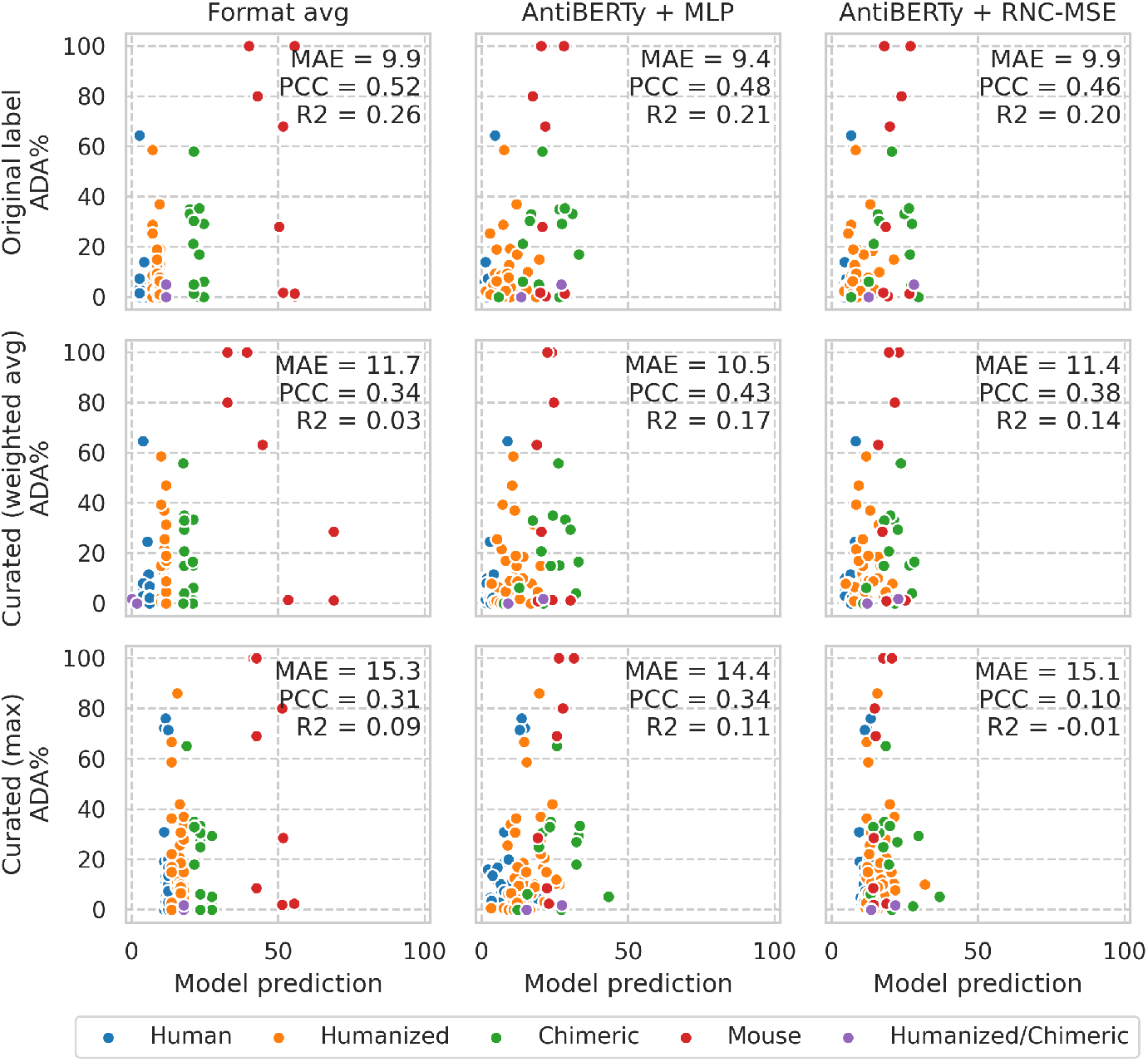
Predicted against observed ADA% for the 110 curated therapeutics (5-folds pooled, mean prediction over 10 runs), organized by model (columns) and label definition (rows), and colored by antibody format. The format lookup’s per-category predictions (which differ slightly in each of the 5 CV splits) align with the original labels but less with the recomputed ones.

**Figure 8:**
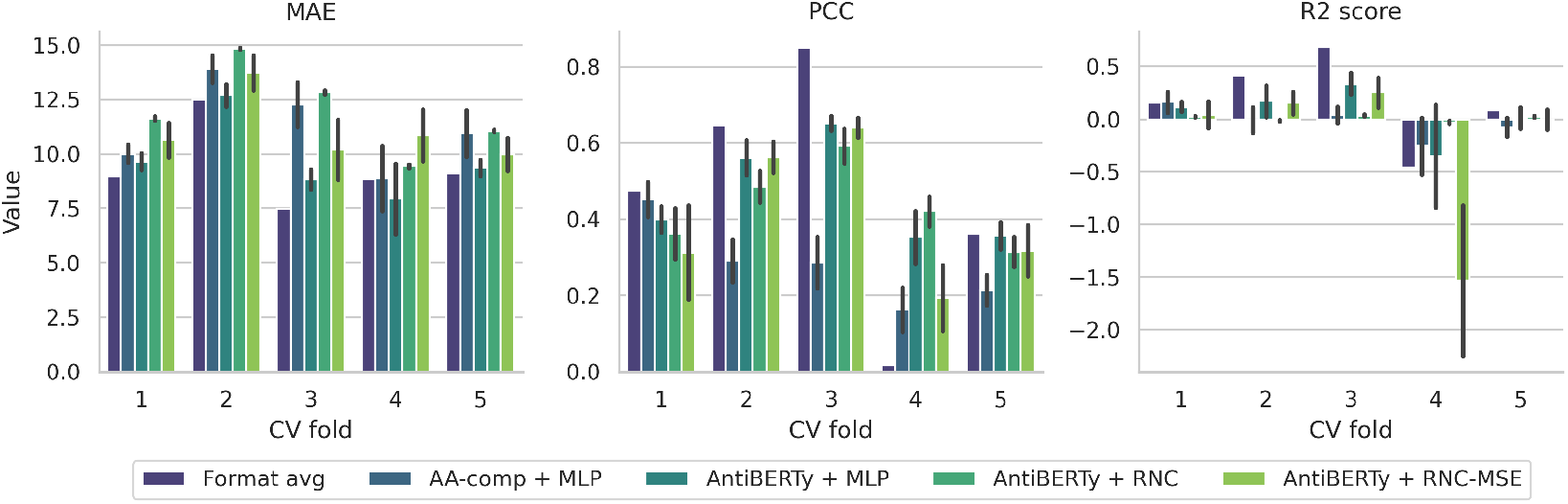
Per-fold MAE, PCC, and *R*^2^ on the 231-antibody dataset (mean *±* SD over 10 runs), for the format lookup and the four learned configurations. Accuracy follows the format composition shown in the cross-validation splits (Figure 5).

**Figure 9:**
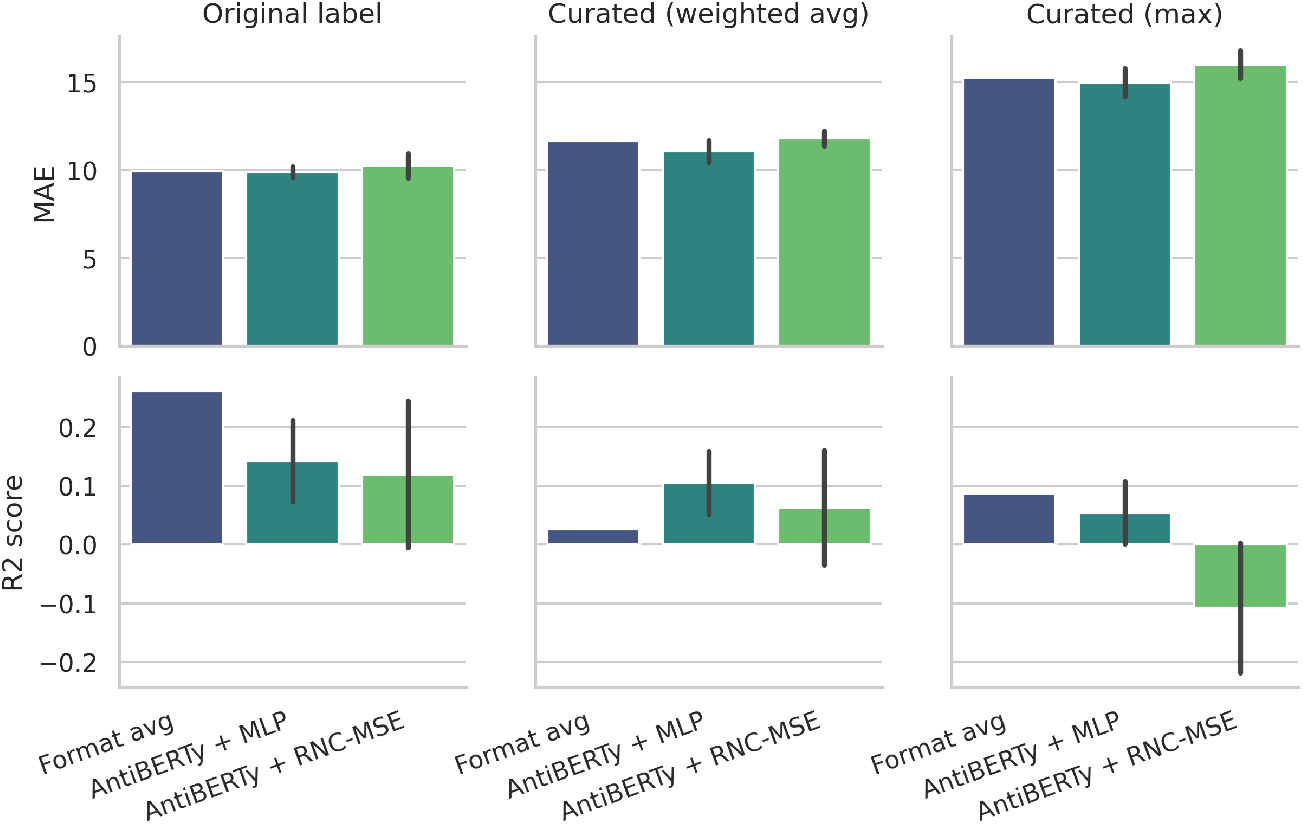
MAE (top) and *R*^2^ (bottom) on the 110 curated therapeutics under the three label definitions (5-folds pooled, mean *±* SD over 10 runs), complementing the PCC results in Figure 4.

**Figure 10:**
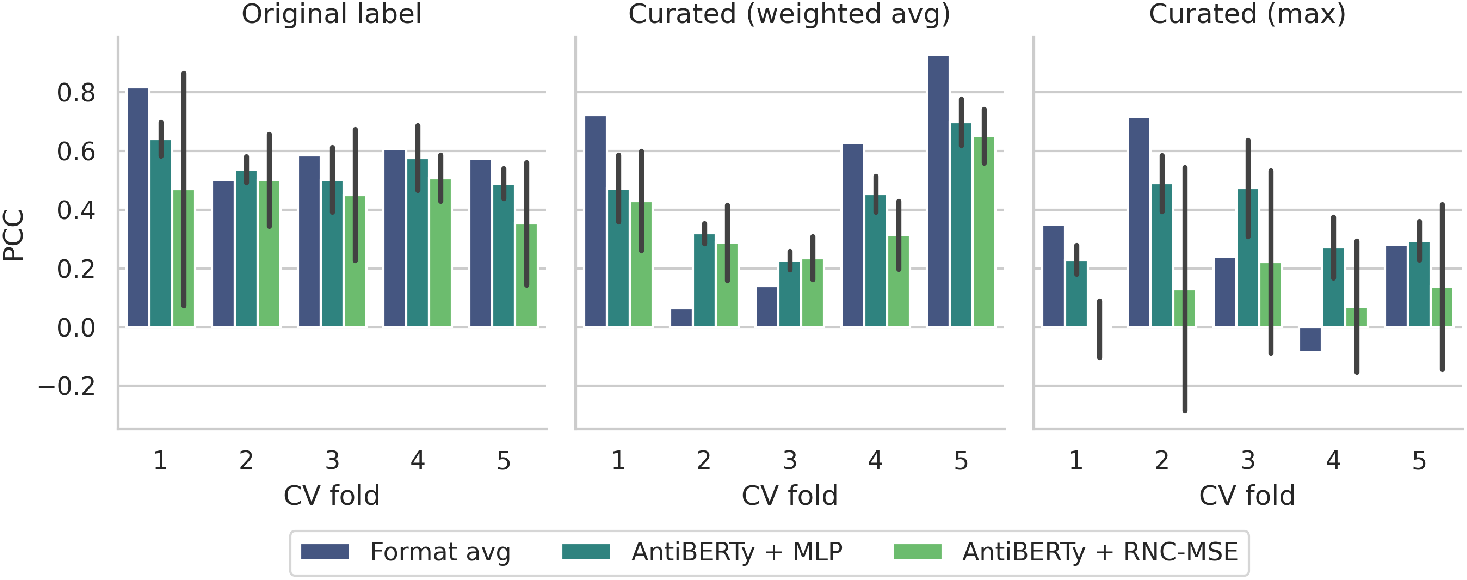
Per-fold PCC on the 110 curated therapeutics under the three label definitions (mean *±*SD over 10 runs). The format lookup’s advantage under the original labels is present in four of five folds and no longer holds consistently under the curated labels, where per-fold estimates vary widely.

## References

[1] Nathaniel R Bennett, Joseph L Watson, Robert J Ragotte, Andrew J Borst, DéJenaé L See, Connor Weidle, Riti Biswas, Yutong Yu, Ellen L Shrock, Russell Ault, et al. Atomically accurate de novo design of antibodies with rfdiffusion. Nature, 649(8095):183–193, 2026.

[2] Jisun Kim, Matthew McFee, Qiao Fang, Osama Abdin, and Philip M Kim. Computational and artificial intelligence-based methods for antibody development. Trends in pharmacological sciences, 44(3):175–189, 2023.

[3] Weijie Zhang, Hao Wang, Nan Feng, Yifeng Li, Jijie Gu, and Zhuozhi Wang. Developability assessment at early-stage discovery to enable development of antibody-derived therapeutics. Antibody Therapeutics, 6(1):13–29, 2023.

[4] Jackie Rao, Ferran Gonzalez, Leon Gerard, and Alexandra Gessner. BOAT: Navigating the sea of in silico predictors for antibody design via multi-objective bayesian optimization. In The 29th International Conference on Artificial Intelligence and Statistics, 2026. URL https://openreview.net/forum?id=eqah46PWof.

[5] Alireza Ghafarollahi and Markus J Buehler. Protagents: protein discovery via large language model multi-agent collaborations combining physics and machine learning. Digital Discovery, 3(7):1389–1409, 2024.

[6] Anna Vaisman-Mentesh, Matias Gutierrez-Gonzalez, Brandon J DeKosky, and Yariv Wine. The molecular mechanisms that underlie the immune biology of anti-drug antibody formation following treatment with monoclonal antibodies. Frontiers in immunology, 11:1951, 2020.

[7] Kathleen P Pratt. Anti-drug antibodies: emerging approaches to predict, reduce or reverse biotherapeutic immunogenicity. Antibodies, 7(2):19, 2018.

[8] Matthew IJ Raybould, Claire Marks, Alan P Lewis, Jiye Shi, Alexander Bujotzek, Bruck Taddese, and Charlotte M Deane. Thera-SAbDab: the therapeutic structural antibody database. Nucleic acids research, 48(D1):D383–D388, 2020.

[9] Taciana Manso, Anjana Kushwaha, Nika Abdollahi, Patrice Duroux, Veronique Giudicelli, and Sofia Kossida. Mechanisms of action of monoclonal antibodies in oncology integrated in IMGT/mAb-DB. Frontiers in immunology, 14:1129323, 2023.

[10] Gemma L Gordon, Matthew IJ Raybould, Ashley Wong, and Charlotte M Deane. Prospects for the computational humanization of antibodies and nanobodies. Frontiers in Immunology, 15:1399438, 2024.

[11] Claire Marks, Alissa M Hummer, Mark Chin, and Charlotte M Deane. Humanization of antibodies using a machine learning approach on large-scale repertoire data. Bioinformatics, 37 (22):4041–4047, 2021.

[12] David Prihoda, Jad Maamary, Andrew Waight, Veronica Juan, Laurence Fayadat-Dilman, Daniel Svozil, and Danny A Bitton. BioPhi: A platform for antibody design, humanization, and humanness evaluation based on natural antibody repertoires and deep learning. mAbs, 14(1):2020203, 2022.

[13] Andrew M Wollacott, Chonghua Xue, Qiuyuan Qin, June Hua, Tanggis Bohnuud, Karthik Viswanathan, and Vijaya B Kolachalama. Quantifying the nativeness of antibody sequences using long short-term memory networks. Protein Engineering, Design and Selection, 32(7): 347–354, 2019.

[14] Aubin Ramon, Montader Ali, Misha Atkinson, Alessio Saturnino, Kieran Didi, Cristina Visentin, Stefano Ricagno, Xing Xu, Matthew Greenig, and Pietro Sormanni. Assessing antibody and nanobody nativeness for hit selection and humanization with AbNatiV. Nature Machine Intelligence, 6(1):74–91, 2024.

[15] Lewis Chinery, Jeliazko R Jeliazkov, and Charlotte M Deane. Humatch-fast, gene-specific joint humanisation of antibody heavy and light chains. mAbs, 16(1):2434121, 2024.

[16] Ariel Tennenhouse, Lev Khmelnitsky, Razi Khalaila, Noa Yeshaya, Ashish Noronha, Moshit Lindzen, Emily K Makowski, Ira Zaretsky, Yael Fridmann Sirkis, Yael Galon-Wolfenson, et al. Computational optimization of antibody humanness and stability by systematic energy-based ranking. Nature biomedical engineering, 8(1):30–44, 2024.

[17] Cade W Gordon, Aniruddh Raghu, Peyton Greenside, and Hunter Elliott. Generative humanization for therapeutic antibodies. In ICLR 2024 Workshop on Generative and Experimental Perspectives for Biomolecular Design, 2024. URL https://openreview.net/forum?id=LiQUkaawXI.

[18] Jian Ma, Fandi Wu, Tingyang Xu, Shaoyong Xu, Wei Liu, Divin Yan, Qifeng Bai, and Jianhua Yao. An adaptive autoregressive diffusion approach to design active humanized antibody and nanobody. bioRxiv, pages 2024–10, 2024.

[19] Fiona A Harding, Marcia M Stickler, Jennifer Razo, and Robert DuBridge. The immunogenicity of humanized and fully human antibodies. mAbs, 2(3):256–265, 2010.

[20] Jonathan Parkinson and Wei Wang. For antibody sequence generative modeling, mixture models may be all you need. Bioinformatics, 40(5):btae278, 2024.

[21] William John Thrift, Jason Perera, Sivan Cohen, Nicolas W Lounsbury, Hem R Gurung, Christopher M Rose, Jieming Chen, Suchit Jhunjhunwala, and Kai Liu. Graph-pMHC: graph neural network approach to MHC class II peptide presentation and antibody immunogenicity. Briefings in Bioinformatics, 25(3):bbae123, 2024.

[22] Shide Liang and Chi Zhang. Prediction of immunogenicity for humanized and full human therapeutic antibodies. PLoS One, 15(8):e0238150, 2020.

[23] Hong Wang, Xiaohu Hao, Yuzhuo He, and Long Fan. AbImmPred: An immunogenicity prediction method for therapeutic antibodies using AntiBERTy-based sequence features. PLoS One, 19(2):e0296737, 2024.

[24] Qiaojing Huang, Yi He, and Kai Liu. Antibody immunogenicity prediction and optimization with ImmunoSeq. bioRxiv, pages 2025–08, 2025.

[25] Zicheng Hu, Sivan Cohen, and Steven J Swanson. The immunogenicity of human-origin therapeutic antibodies are associated with V gene usage. Frontiers in Immunology, 14:1237754, 2023.

[26] Pawel Dudzic and Konrad Krawczyk. Herding cats: predicting immunogenicity from heterogeneous clinical trials data. bioRxiv, pages 2025–10, 2025.

[27] Michael Chungyoun and Jeffrey Gray. Fitness landscape for antibodies 2: Benchmarking reveals that protein AI models cannot yet consistently predict developability properties. bioRxiv, 2025.

[28] Chantal T Harris and Sivan Cohen. Reducing immunogenicity by design: Approaches to minimize immunogenicity of monoclonal antibodies. BioDrugs, 38(2):205–226, 2024.

[29] Jeffrey A Ruffolo, Jeffrey J Gray, and Jeremias Sulam. Deciphering antibody affinity maturation with language models and weakly supervised learning. Machine Learning for Structural Biology Workshop, NeurIPS, 2021.

[30] Kaiwen Zha, Peng Cao, Jeany Son, Yuzhe Yang, and Dina Katabi. Rank-N-Contrast: Learning continuous representations for regression. Advances in Neural Information Processing Systems, 36:17882–17903, 2023.

[31] Oliver M Turnbull, Dino Oglic, Rebecca Croasdale-Wood, and Charlotte M Deane. p-iggen: a paired antibody generative language model. Bioinformatics, 40(11):btae659, 2024.

[32] Sudhanshu Agnihotri, Bruno Gonzalez-Nolasco, Brinda Monian, Sofie Pattijn, Chloé Ackaert Patrick Wu, Hubert Kettenberger, Sophie Tourdot, Timothy Hickling, Zicheng Hu, et al. The immunogenicity database collaborative: a standardized, publicly available database for clinical immunogenicity observations and insights. Frontiers in Immunology, 17:1816949, 2026.

